# Integrated host–phage RNA seq analysis of Mycobacteriophage D29 infection reveals dual-arm balanced phage genome expression and inhibited expression of host transporter and VapC target genes

**DOI:** 10.1101/2025.11.21.689395

**Authors:** Rahul Shaw, Shrestha Ghosh, Subhajit Dutta, Anirban Ghosh, Sujoy K. Das Gupta

**Author notes:** Dept. of Zoology, Government Navin College Odgi, District- Surajpur, Chhattisgarh, Pin code 497231, India (Rahul Shaw). Department of Microbiology, Tumor and Cell Biology (MTC), Karolinska Institutet, Biomedicum C9, Solnavägen 9, 171 65 Solna, Sweden. Department of Transfusion Medicine, All India Institute of Medical Sciences, Bhubaneswar India (Anirban Ghosh). To whom correspondence should be addressed. Sujoy K. Das Gupta.

## Abstract

The Mycobacteriophage D29 infects Mycobacterium species. Many of these are deadly human pathogens. In earlier studies, we performed proteomic analyses using a temperature-sensitive variant of D29, called D29ts12. This work indicated that a regulatory mechanism controls the expression of the phage’s right arm genes. In this study, we executed a transcriptomic analysis. We gained further insight into how phage infection affects gene expression, both in its genome and in the host genome. We identified several novel non-coding RNA contigs spanning the cos site junction. This finding indicates that this region is expressed. In addition, we observed two areas of intense transcriptional activity. One is located downstream of P_left_, in the right arm, and coincides with a major noncoding RNA species. This RNA species is essential for lytic growth, as reported earlier for mycobacteriophages of the A2 family. The other area of transcriptional activity was gene 17, which encodes the major coat protein. By comparing TPM values, we found that thermal inactivation of phage D29ts12 growth significantly decreased right-arm gene expression. Interestingly, an accompanying sharp increase in left-arm gene expression was observed. Therefore, a mechanism ensures balanced expression from the two arms. We also analysed the impact of phage infection on host gene expression. The results indicate significant changes in the expression levels of genes encoding transporters and genes regulated by the toxin VapC. This investigation is the first to use a mycobacteriophage to examine the combined transcriptome of the phage and its host.

## INTRODUCTION

Mycobacteriophages are regarded as toolboxes for TB research (1). Like other bacteria, *E. coli,* for example, these phages have been extremely useful in genetically manipulating mycobacteria. However, in addition to their use in genetic engineering, mycobacteria have been studied extensively to understand the evolution of phage genomes. Ongoing efforts by Graham Hatfull’s lab have led to the discovery and sequencing of a vast repertoire of mycobacteriophages. Interestingly, many of these phages were isolated by school and graduate students as part of an educational exercise (2). These phages are likely to be useful for various reasons, including phage therapy. Cocktails of mycobacteriophages have been used to treat patients suffering from TB caused by *Mycobacterium abscessus.* (3).

The ability of phages to lyse their host cells is fundamental to their existence. However, host cell killing is not necessarily only due to lysis. Even without lysis, a bacterial population can die in the presence of a phage. The phenomenon is known as abortive infection, in which the host cell commits suicide, preventing the phage from multiplying and spreading (4). Recent studies indicate that host-encoded anti-phage systems, such as CBASS, can contribute to host cell defence mechanisms (5). This investigation is based on an earlier study from this lab that demonstrated that mycobacteriophage D29 causes cell death without lysis (6). This inference was drawn from the observation that the medium’s optical density decreased only marginally, even though most cells were rendered non-viable. According to the study, a model was proposed, indicating that mycobacteriophage D29 may induce cell death through multiple mechanisms. It is becoming exceedingly clear that phage-infected cells undergo a phenomenon similar to apoptosis. Self-killing of the infected bacterium minimises the further spread of the phage (7). A similar phenomenon occurs with phage D29(8).

Phage D29 is a well-accepted model for studying how mycobacteriophages infect mycobacteria(9). It belongs to the A2 family of mycobacteriophages, including L5, a closely related member (10). In the genomes of D29-related phages, gene expression takes place in a convergent manner from the two ends of the genome. The open reading frames are organised in two transcriptional units, one running leftwards from the right end (right arm) and the other in the opposite direction from the left (left arm)(11). In the right arm, multiple genes code for proteins involved in nucleotide metabolism (12). The left arm, on the other hand, harbours genes that code for structural proteins, mainly the capsid proteins and those involved in cell lysis(13). The consensus is that genes on the right are expressed early, whereas those on the left are expressed late. However, there is little evidence to suggest that a mechanism exists by which the phage switches gene expression from the right to the left. The genome of L5, a closely related phage, has cohesive ends (14). The same cohesive sequence is also present in phage D29(11). Therefore, both L5 and D29 are likely circularised following infection; however, there is no evidence that the junction region is also expressed.

Transcriptomic studies have been performed earlier using D29 and several other phages (15). These studies revealed that, early in infection, the right-arm genes are expressed first, followed by the left-arm genes. Gene expression appears to be regulated not only by promoters located at the extreme ends of the genome but also by internal elements that overlap with the operator-like motifs known as stoperators (16, 17).

The primary issue that remains unsolved is the mechanism by which the shift from the left to the right arm occurs. Does a molecular switch exist? To answer this question, it is necessary to develop phage D29 mutants that are conditionally defective in gene expression regulation. We previously reported a temperature-sensitive mutant, D29ts12, that cannot grow at the non-permissive temperature of 42 °C, but grows typically at 37 °C. Using this mutant, we performed proteomic studies. From such studies, it became apparent that a positively acting factor may be involved in the regulation of the right arm genes. (18). However, the proteomic analysis did not give a detailed picture of every gene, particularly those that are less expressed. We therefore proceeded to perform transcriptomic analysis to obtain a deeper insight into how phage D29 gene expression is controlled.

In our previous studies, we attempted to analyse what happens to the host during phage infection. Electron Microscopy and other cell biological studies revealed significant changes in the host *Mycobacterium smegmatis* cell following mycobacteriophage gene expression (8). One striking result was a significant increase in VapC RNA levels compared with VapB. Since VapC is known to have RNase activity (19) therefore, one would expect that RNAs derived from the host genome that are VapC substrates will be degraded following D29 infection. Since we anticipated changes in the host expression pattern following phage infection, we have also analysed the host transcriptome alongside the phage transcriptome. In this study, we present a detailed transcriptomic analysis of *M. smegmatis* cells infected with D29.

## MATERIALS AND METHODS

### Bacterial strains, phages, culture and infection conditions

*M. smegmatis* mc^2^155 (Msm) cells were grown in Middlebrook 7H9 medium (20) (Difco) with 0.01% Tween-80 and 0.2% glycerol. D29 phage suspensions were prepared by confluent lysis. D29 phage infections were performed using a log phase culture of Msm at a multiplicity of infection of 1 in the presence of 2mM CaCl2 as described previously (20). For plaque formation, the hard agar was overlaid with top agar with 2 mM CaCl_2_.

### Chemicals and reagents

Restriction enzymes, Taq DNA polymerase, and other DNA-modifying enzymes were obtained from New England Biolabs or Thermo Scientific. All other chemicals of the highest purity grade were procured from SRL, MERCK, or Sigma-Aldrich. DNA oligonucleotides were obtained from Eurofins.

### RNA isolation

RNA isolation was performed by the Trizol method. 20 ml of Msm culture, phage-infected or uninfected, was centrifuged for 5 min at 6000 g. The pellet was resuspended in 2 ml of TE buffer containing lysozyme (5 mg/ml) and incubated at 37°C for 30 minutes. Trizol (5ml) was then added, and the mixture was stirred thoroughly using a pipette. One ml of chloroform was then added, followed by vigorous shaking. The phases were separated by centrifugation for 20 min. at 13000 g. The aqueous phase was transferred to a Corex tube, and 0.7 volumes of cold isopropanol were added to precipitate the RNA. The sample was incubated at room temperature for 30 min. followed by centrifugation at 13000 g for 15 min. The pellet obtained was washed with 70% ethanol. The pellet was dispersed in 30 µL of DEPC-treated water, air-dried for 20–30 min, and then incubated at 55°C for 10 min. Single-use aliquots of RNA were kept at -20°C for short-term use or at -80°C for long-term use. The RNA quality and quantity was measured using Nanodrop, UV-spectrophotometry.

### RNA-seq analysis

The extracted RNA samples were outsourced for RNA-seq analysis. In brief, the submitted RNA was depleted with the Ribocop rRNA depletion kit according to the manufacturer’s protocol (Lexogen #125–127, Greenland, NH, USA). In total, 100 ng of ribodepleted RNA was utilised and converted into the final library using the TruSeq stranded RNA library preparation kit (Illumina, USA) following the manufacturer’s protocol.

### Contig analysis

Raw sequencing reads from four transcriptomic datasets were preprocessed to remove adapter sequences and low-quality bases using Trim Galore v0.6.6 with default parameters. Reads with Phred scores below 20 were discarded. The resulting high-quality reads (Q20-filtered) were aligned to the genome of Mycobacterium virus D29 (NCBI Reference Sequence: NC_001900.2) using Bowtie2 v2.5.1 under default settings(21). The genome-aligned reads from each dataset were independently assembled using SPAdes v3.13.1 in rnaSPAdes mode(22). The resulting contigs from each dataset were mapped back to the D29 genome using Bowtie2 v2.5.1 with default parameters. Alignment outputs were subsequently analyzed using Samtools v1.13(23). CDS-aligned reads were used to characterise RNAs associated with coding regions, while CDS-unaligned reads provided insights into RNAs potentially originating from non-coding regions. The final genome-aligned contigs were utilised to determine the genomic positions of long transcript stretches, facilitating structural and functional annotation of the D29 transcriptome.

### Differential gene expression

For differential gene expression calculations, the DESeq2 tool was used. The final statistical analysis was performed, and TPM (transcripts per million) values were determined. The statistical significance was expressed as P values. The reference genomes used for TPM calculation were mycobacteriophage D29 and that of *Mycobacterium smegmatis* mc^2^155 (NC_008596.1). The sequence data are available in the SRA databases under Bioproject PRJNA1153798.

Two different strategies were used: a) Infection of Msm with D29ts12, a mutant of phage D29, that grows at 37°C, but not at 42°C. The RNA was isolated from D29-infected cells grown at 37°C or 42°C for 30 or 120 minutes. The phage transcriptome derived from the 37°C (30 mins.) sample was compared with that of 42°C (30 mins.) and, similarly, 37°C (120 mins.) with that of 42°C (120 mins.). For P-value consideration, the host transcriptome was analysed using the same temperature samples as duplicates. This approach was taken as it was found that the intensities of the 37°C _30 min. and 37°C _120 min. pairs were well-correlated, and the same was true for the 42°C pair (Fig. S1). (b) Isolation of RNA from cells either infected or uninfected (control) with the wild-type D29 at two time points, 30 min and 120 min. There were four samples in this experiment: two uninfected (30 mins. and 120 mins.) and two infected samples (30 mins. and 120 mins.). Although both sets were sequenced, we focused on the 120-min. set (uninfected and infected). The 120-minute combination was analysed by RT-PCR and NGS analysis.

## RESULTS

### Novel transcripts from the D29 phage genome were identified through contig analysis

To investigate the regulation of gene expression from phage D29ts12 in more detail, we performed NGS analysis of RNA extracted from phage-infected cells at two time points (30 min and 120 min) and two temperatures (37 °C and 42 °C). The total RNA includes both phage-derived RNA and the host. We first performed contig analysis for the phage genome using the sample from the 37 °C, 120 min time point. At this temperature and time interval, transcriptional activity from the phage genome is at its maximum. The results show that almost 99% of the genome was covered by the NGS reads (Fig. 1A, red bars). Only a small region spanning ORFs 72 and 73 remained uncovered. Interestingly, we found that the sequence corresponding to the junction of the two arms (the cos site) is also expressed; hence, we conclude that the genome is circularly permuted (Fig. 1B).

**Figure 1.**
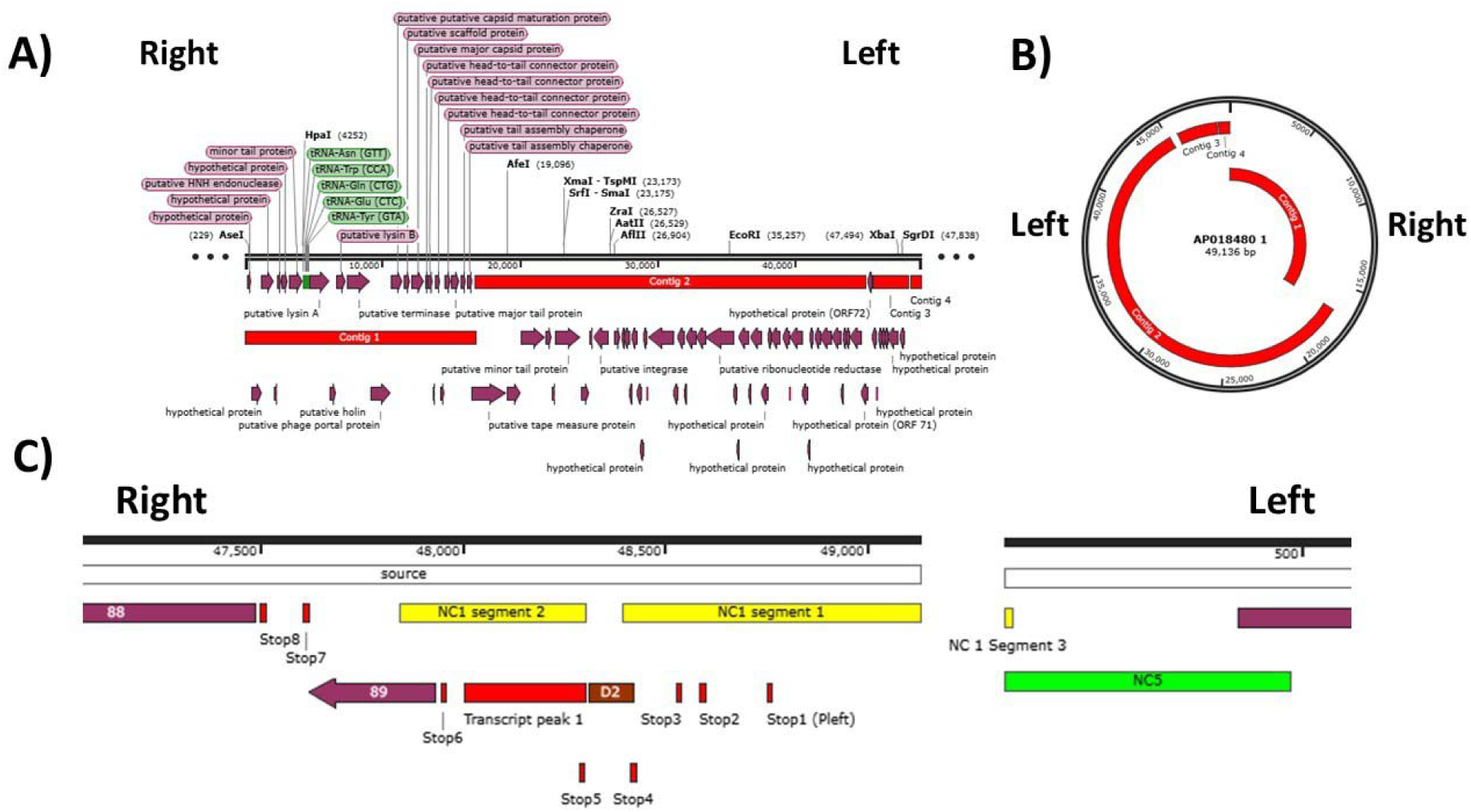
Schematic representation of the Phage D29 contig map obtained from RNA seq analysis of D29-infected cells, drawn using Snapgene software (A). A simplified circular map is shown in (B). The red bars or arcs represent regions covered by the reads obtained. (C) Detailed analysis of the junction between the left and right arms. The red vertical bars represent stoperator sequences. The first stoperator (Stop 1) overlaps with P_left_. Non-coding regions are indicated as NCs. There are three yellow boxes labelled NC1 segments 1, 2, and 3. These three regions together constitute a single contig. The gap between 1 and 2 represents a deletion in the strain (box D2) used in this study relative to the wild type. Segment 3 is located on the other side of the cos site. If the two arms are joined at the cos site, then this contig (NC1) will span approximately 1.2 kb. The red colored box below NC1 segment 2 represents a region that is highly expressed (refer to Fig. 2, peak on the right arm). The green segment NC5 is also a novel contig identified in this study. Since RNA-seq does not provide strand information, the directionality could be either orientation.

To gain a deeper understanding of the transcriptome in the cos site region, we created a detailed map for this region. We identified a major contig, 1169 nucleotides in length, spanning the cos site (Fig. 1C). The yellow blocks represent this contig, located on the extreme right, specifically NC1 segments 1 and 2 of the right arm, and NC1 segment 3, a small segment on the extreme left of the left arm. NC1 segments 1 and 2 are continuous but are depicted as bipartite because a 111-bp deletion (D2) is present in this region relative to D29 (AF02221) in the D29 strain used. If we join all the yellow segments, we will obtain a contig (NC1) spanning the left-right junction. At the same time, we observed another contig, NC5, that partly overlaps NC1 segment 3 at the extreme left end of the left arm. The results indicate that the replicative form of the genome is circular and that the junction between the left and right ends is transcribed.

### Raw read alignment maps indicate two intensely transcribed regions in each arm

The raw results indicate that the entire genome is heavily transcribed, except for a region located at the junction of the two convergent operons spanning gene 33 (the integrase). Peak activities were observed in two areas. On the right arm, a sharp peak was observed immediately upstream of gene 89 (Fig. 2, Peak 1, Fig. 1C red box), whereas on the left, a peak (Peak 2) was observed corresponding to gene 17 (major capsid protein) (Fig. 2, indicated by red arrows). The sharp peak on the right arm appears to conform to the leftmost non-coding RNA peak reported earlier, which was found to be essential for lytic growth (15). The results show that when comparing the 37 °C 30-minute and 120-minute time points, the observed patterns are very similar, though intensities vary. If we now compare the 42 °C results with those at 37 °C, we find that at the non-permissive temperature, expression from the left arm decreases significantly at both time points. The intensity of peak 2 on the right decreases and is barely visible. The results are consistent with those of our earlier proteomic study(18).

**Figure 2.**
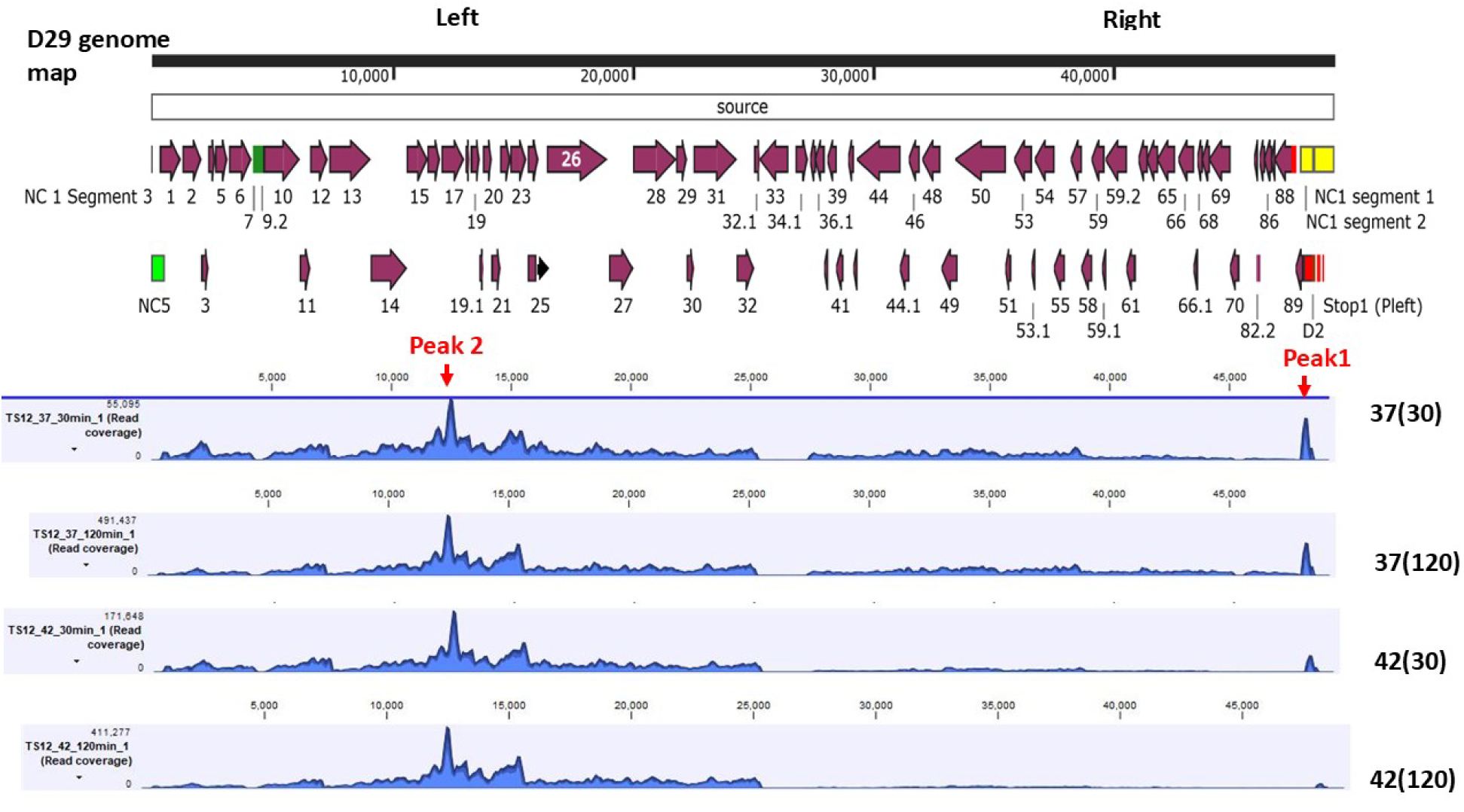
Raw read coverage maps. The transcriptomes derived from Msm cells infected by D29ts_12, at either 37°C (30 or 120 min.) or 42°C(30 or 120 min.) as indicated, 37(30), 37(120), 42(30), 42(120) were analyzed for phage-specific transcripts. The maps obtained were manually aligned with the D29 genome map, showing the positions of the various genes and the cos site-derived novel non-coding contigs (NC1 segments 1-3, NC5; yellow and green boxes, respectively). Since the alignment was done manually, the correspondence is approximate. The red arrows point towards two peaks of intense activity in the map.

### Normalised expression (TPM) data indicate a novel control mechanism that balances expression from the two arms

Conversion of the raw read data to data normalised by gene size and sequencing depth provided further insight into how gene expression from the phage genome was controlled. In both cases, left-arm genes are more expressed than right-arm genes. However, as expected, the level of expression of the phage genes was marginally higher at 120 mins (Figure 3D compared to C). At 42°C, remarkable changes in the phage transcriptome were observed relative to 37°C (Fig. 3F compared to E). The right-arm gene transcripts (Fig. 3F, boxed region) were significantly reduced in intensity, as expected. But, at the same time, the left arm gene transcripts were remarkably elevated. This unexpected phenomenon is more clearly observed when the fold change in gene expression is examined for each phage gene as the temperature is shifted from 37°C to 42 °C (Figs. 3G and H for 30 and 120 min., respectively). If the results from 30 and 120 minutes are considered biological replicates, it is possible to create a volcano plot that accounts for p-values. The volcano plot results are consistent with the observations above (Fig. 3I). There is a significant decrease in right-arm transcripts and, at the same time, an increase in left-arm transcripts. This result suggests that a D29-coded factor, temperature-sensitive in the D29ts12 mutant, ensures balanced expression from both arms of the phage genome. The factor, when active, appears to positively influence the expression of right-arm genes and negatively influence the expression of left-arm genes. Conversely, its inactivity causes reduced expression in the right arm and increased expression in the left.

**Figure 3.**
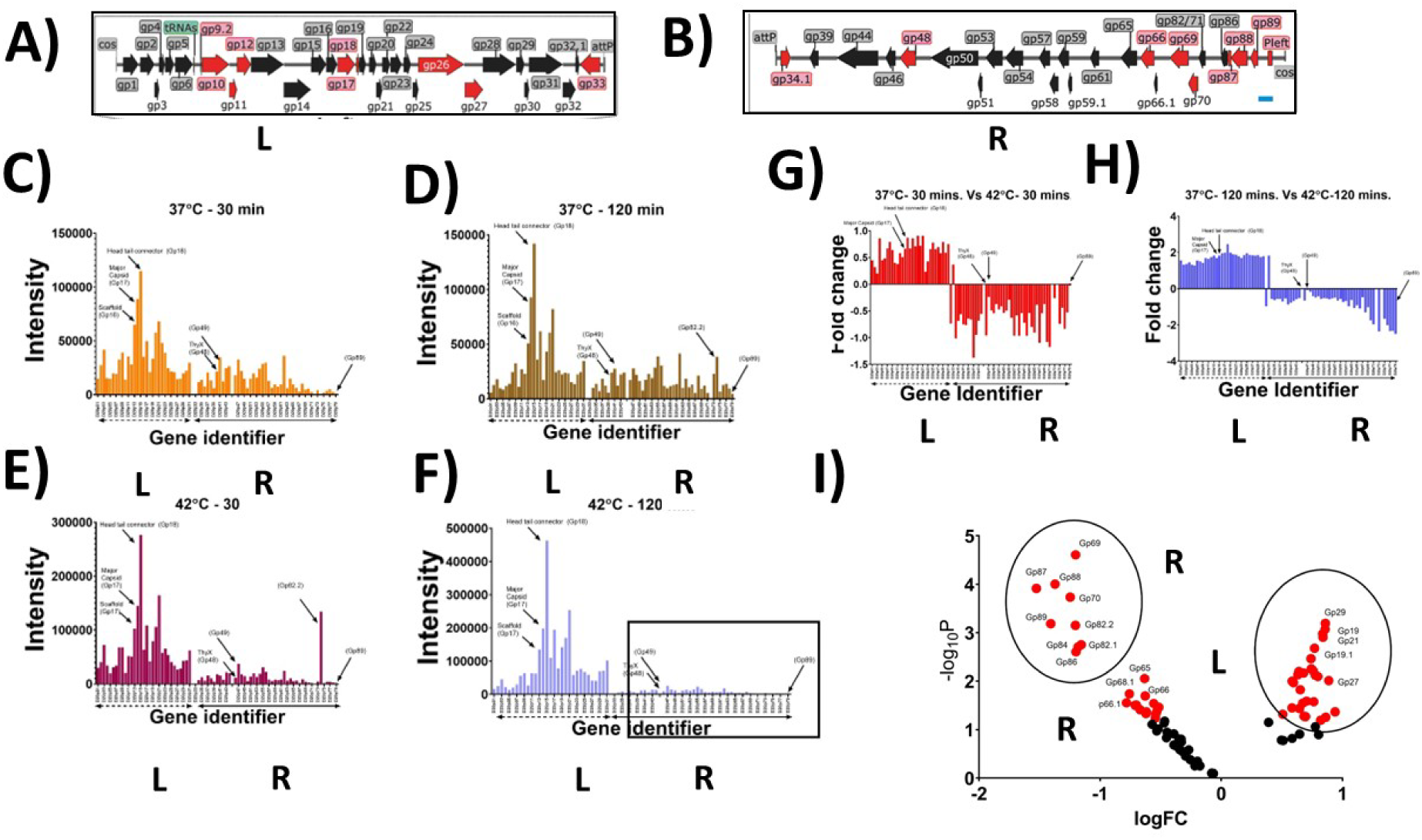
RNA-seq analysis of the D29-infected *Mycobacterium smegmatis* transcriptome. A) and B) Schematic representations of the Left (L) and Right arms (R) of phage D29, split in the middle at the attP site. C) to F) Intensity (TPM) of phage gene expression after 30 or 120 min infection as mentioned. Fold changes in gene expression observed upon shifting from 37 °C to 42 °C are shown for 30 min (G) and 120 min (H). A volcano plot (Log Fold Change vs Log of P values) was derived (I) by treating 30 and 120 min as duplicates.

### Effect of Phage Infection on Host Gene Expression

All four RNAseq results were then analysed to investigate the effect of phage infection on host gene expression. The phage transcript intensities of the 37°C _30 min. and 37°C _120 min pair were well-correlated, and the same was true for the 42°C pair (Fig. S1). Therefore, while analysing the host transcriptome, we considered 37 °C _30 min and 37 °C _120 min as one pair and 42 °C _30 min and 42 °C _120 min as another pair. The average expression of host genes derived from the 42 °C pair was then compared to that of the 37 °C pair. The results of this analysis indicated that 132 genes were significantly (p-value < 0.05) down-regulated at 42 °C compared to 37 °C, whereas 81 were up-regulated at 42 °C compared to 37 °C.

The pathways likely to be affected differentially were then analysed using KEGG pathway analysis. A summary of the results obtained is presented in Tables 1 and S1. The results indicated that the most significantly affected genes (Enrichment FDR, 1.6 x E-05) belong to the categories of ABC transporters and metabolic pathways. Because the members of the metabolic pathways were highly diverse, we focused on the well-defined ABC transporter category.

**Table 1.** Functional enrichment of *M. smegmatis* genes differentially expressed following D29 phage infection.

| Functional category | Representative functions/pathways | No. of genes | Example gene IDs | Enrichment FDR |
| --- | --- | --- | --- | --- |
| **Transport and membrane systems** | ABC transporters (peptide, metal ion, lipid uptake/export) | 11 | *MSMEG_0012*,<br>*MSMEG_0015*,<br>*MSMEG_5781* | $1.6 \times 10^{-5}$ |
| **Central and secondary metabolism** | Metabolic pathways (amino acid, carbohydrate, cofactor biosynthesis) | 23 | *MSMEG_0911*,<br>*MSMEG_3043*,<br>*MSMEG_3951* | $7.2 \times 10^{-4}$ |
| **Virulence-associated pathways** | Homologs of *M. tuberculosis* genes linked to stress and persistence | 3 | *MSMEG_0709*,<br>*MSMEG_1583*,<br>*MSMEG_5782* | $1.4 \times 10^{-3}$ |
| **Motility and signalling** | Bacterial chemotaxis, two-component signalling | 2 | *MSMEG_3598*,<br>*MSMEG_3599* | $9.5 \times 10^{-3}$ |
| **Lipid metabolism** | Fatty acid biosynthesis and degradation, branched-chain amino acid metabolism | 7 | *MSMEG_0655*,<br>*MSMEG_3951*,<br>*MSMEG_4717* | $2.0 \times 10^{-2}$ |
| **Sulfur and selenium metabolism** | Sulfur assimilation and selenoprotein synthesis | 5 | *MSMEG_4977–4979* | $2.1 \times 10^{-2}$ |
| **Nucleic acid processes** | RNA degradation and DNA replication | 4 | *MSMEG_0709*,<br>*MSMEG_6275* | $2.1 \times 10^{-2}$ |
| **Energy metabolism** | Pyruvate and pyrimidine metabolism | 6 | *MSMEG_3043*,<br>*MSMEG_4711*,<br>*MSMEG_4712* | 0.027–0.049 |

We listed the genes under the ABC transporter category, along with the functions of the proteins they encode (Table 2). We can categorise the differentially accepted genes into the following: a) Ferric-enterobactin transport (Fep system), b) Phosphonate / Polyamine transport, c) phosphate transfer, d) Sugar transport/regulation, e) hydrophobic amino acid transporter. Except for the sugar transport category, the expression levels of the others decreased at 42 °C compared with 37 °C. We suspected that the differential expression of the host gene may be an indirect consequence of the temperature shift’s effect on viral genome expression. However, the differential effects could also be due to heat shock. To resolve this issue, we infected the host with the wild-type phage. We examined the expression status of genes representing the above categories using RT-PCR as described in the following section.

**Table 2.** Differential expression of ABC-transporter genes in *Mycobacterium smegmatis* infected with the temperature-sensitive phage D29ts12.

| Functional subclass | Representative gene(s) | Regulation (42 °C vs 37 °C) | Mean Log<sub>2</sub> fold change | Functional comment |
| --- | --- | --- | --- | --- |
| **Ferric-enterobactin transport (Fep system)** | *MSMEG_0012,0015,0018 0020,* | ↓ at 42 °C (↑ at 37 °C) | −0.8 to −1.5 | Iron uptake is suppressed under non-permissive temperature |
| **Phosphonate / Polyamine transport** | *MSMEG_0649, 0659* | ↓ at 42 °C (↑ at 37 °C) | −1.6 to −1.9 | Reduced uptake of phosphonates and polyamines |
| **Phosphate transport (Pst system)** | *MSMEG_5781, 5782* | ↓ at 42 °C (↑ at 37 °C) | −0.7 to −1.4 | Downregulation of phosphate assimilation |
| **Sugar transport/regulation** | *MSMEG_3598, 3599* | ↑ at 42 °C (↓ at 37 °C) | +0.44 to +0.45 | Possible compensatory induction of carbohydrate import and regulation |
| **Hydrophobic amino acid transport** | *MSMEG_6880* | ↓ at 42 °C (↑ at 37 °C) | −0.55 | Indicative of nutrient stress and altered amino acid uptake |

### Transcriptomics of wild-type D29 infection

In the previous section, we used a temperature-sensitive mutant to understand how the phage and host genome expression patterns change upon phage infection. The investigation provided us with specific leads regarding which host genes are affected by alterations in phage gene expression. Does the expression of the same genes change when Msm is infected with wild-type phage D29? To address this issue, we infected Msm with phage D29 and analysed the transcriptome. To verify that infection had occurred, we recovered the infected cells at 120 mins. time point and isolated the RNA. We performed RT-PCR and NGS analysis to determine whether infection had occurred and whether the phage genes were transcribed (Fig. S2). For the RT-PCR test, we targeted gene 88, located on the right arm, which we had previously found to be expressed reproducibly during phage D29 infection (17). Once we were convinced that the phage-specific genes were optimally expressed in the target cells, we analysed the expression status of the transporter genes identified in the previous section as differentially regulated. For this purpose, we targeted at least one gene from each set (Fig. 4). The genes tested were MSMEG_0020 (FxuD, iron utilisation), MSMEG_3598 (sugar binding, periplasmic sugar binding), MSMEG_3599 (sugar binding) and MSMEG_5782 (Psts, Phosphate transfer). The results indicate that the expression of the representative genes is significantly reduced. We also investigated MSMEG_6880, as it encodes a transporter for hydrophobic amino acids (Fig. 4, lower panel). MSMEG_6880 was found to be differentially regulated in the D29ts12 experiment (Table 2). However, MSMEG_6880 is part of an operon encoding proteins involved in amino acid transport. Therefore, we also investigated these genes (Fig. 4E). Our results indicate that the expression of three genes in this operon (MSMEG_6880, 6879, and 6878) was significantly affected, whereas that of MSMEG_6877 was marginally affected. The results suggest that all the genes affected differentially in the experiment with D29ts were downregulated in the experiment with the wild-type D29 phage.

**Figure 4.**
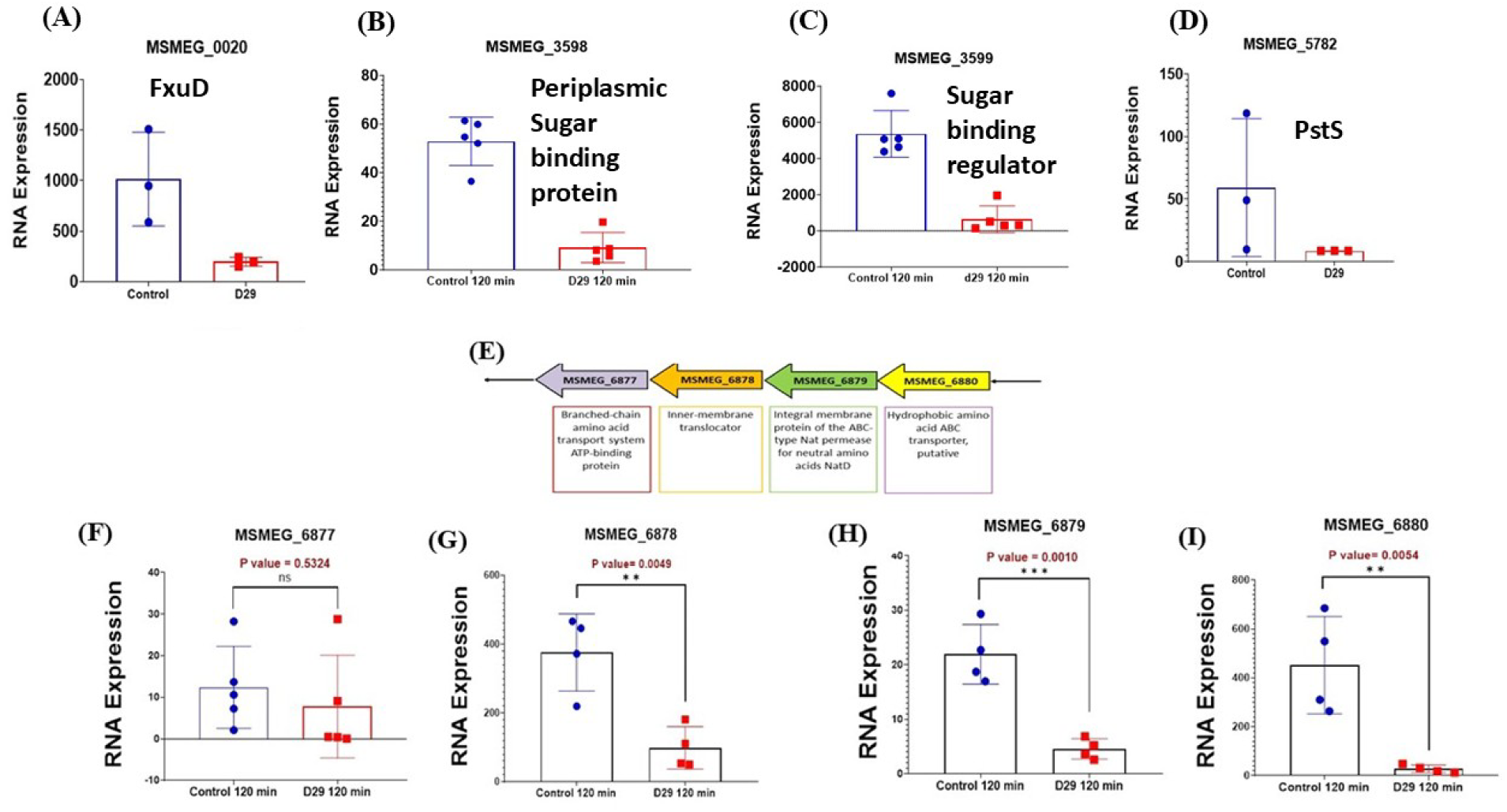
Inhibition of Msm gene expression caused by phage D29 infection. The expression of the host genes that were differentially expressed at 42°C relative to 37°C in D29ts12-infected cells was independently assessed by infecting Msm with wild-type D29, extracting RNA and performing comparative real-time PCR analysis using rRNA as the internal control. The genes tested are indicated. Wherever necessary, significance levels are indicated.

### Effect of phage infection on expression of Carbohydrate metabolism-related genes

The increased activity of VapC in infected Msm cells relative to VapB was an intriguing observation reported in our previous study (24). If this observation were accurate, then it is likely that all transcripts known to be targets of VapC in Msm would be downregulated. In a previous study, the effect of VapC overexpression on the stability of Msm RNA was studied(19). The results showed that overexpression inhibited the expression of 42 genes, all of which encode proteins involved in carbohydrate metabolism. We asked: if phage infection leads to higher VapC activity, then the effect would be the same as overexpressing VapC. Accordingly, we examined the D29 infectome NGS data for differential expression of carbohydrate metabolism-related genes. We found that, excluding three apparent exceptions and a few borderline cases, the expression of 36 of the 42 genes reported in the previous study was adversely affected (Figs. 5A green cluster compared to red and Table S2).

**Figure 5.**
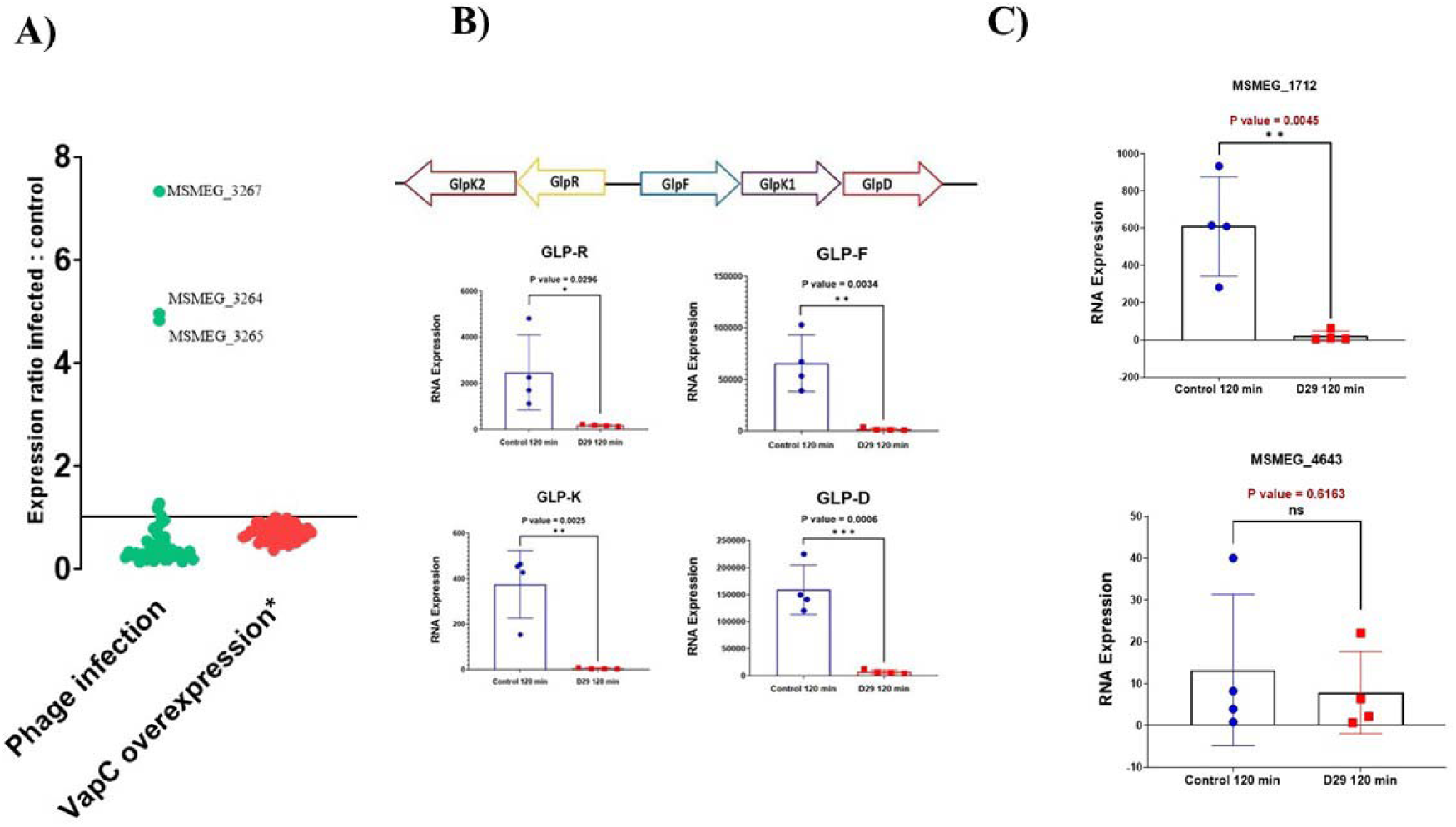
A) Effect of phage infection on the expression of Carbohydrate metabolism-related genes. The results of the VapC overexpression (red cluster) are being reproduced from another study (19) after taking permission from the authors, the detailed results are available in the Supplementry file. B) Effect of phage infection on the expression of glycerol metabolism-related genes and (C) Expression of two marker genes MSMEG_1712 and MSMEG_4643 that are differentially expressed in Msm cells overexpressing VapC relative to VapBC.

The list of 42 genes includes several enzymes involved in glycerol metabolism. We tested how phage infection affects the genes involved in glycerol metabolism. This experiment was performed by doing RT-PCR. The results of such an experiment confirmed that the expression of all the glycerol metabolism-related genes Glp-R, Glp-F, Glp-K and GlpD was significantly diminished (Fig. 5B)

We have also selected two genes differentially regulated in VapC-overexpressing cells as markers of VapC activity. One such gene was MSMEG_1712, the expression of which in VapC-only-expressing cells was downregulated by 6 to 8 fold relative to VapBC-expressing cells (19). The other was MSMEG_4634, which increased moderately by 2- to 3-fold. We argued that if VapC was activated in D29-infected cells, then expression of MSMEG_1712 should decrease relative to uninfected cells, whereas that of MSMEG_4643 should increase. We carried out this experiment using RT-PCR. The results reveal that expression of the marker gene MSMEG_1712 was significantly reduced (Fig. 5C, upper panel). In the case of MSMEG_4643, there was neither an increase (Fig. 5C, lower panel) nor a significant decrease. The downregulation of MSMEG_1712, but not MSMEG_4643, indicates that VapC is activated in D29-infected cells. In the MSMEG_4643, there is a minor deviation. No increase was observed, but most importantly, no decrease happened. This minor deviation may be due to differences in the experimental setups. However, the significant downregulation of MSMEG_1712 provides confidence that the effect of phage D29 is VapC-mediated.

## DISCUSSIONS

In recent years, several groups, including our laboratory, have attempted to understand how the D29 phage genome is expressed (18, 25). To understand how gene expression in the phage D29 is differentially regulated, we generated a temperature-sensitive mutant, D29ts12, and characterised it using growth assays and proteomic analysis. The results indicated that in this mutant, some regulatory function must have become thermolabile, which is why the peptides derived from proteins encoded by the right arm genes became undetectable in our LC-MS-MS analysis experiments (18). Although the results provided some insight into how the right-arm genes were controlled, the low sensitivity of the mass-spectrometry-based approach limited the feasibility of a detailed quantitative interpretation. Therefore, we shifted to NGS sequencing, which enabled us to examine phage D29 gene expression at greater detail. At the same time, we looked at how the host transcriptome is altered following phage infection. We used the temperature-sensitive mutant D29ts12 and the D29 wild-type in this study.

From the results, we identified RNA species corresponding to the entire genome. It is generally accepted that there are two promoter elements in the extreme ends, R_left_ and P_right_, from which transcription of the genome takes place. P_left_ is located between nt positions 48750 and 48762, about 262 bp downstream from the right end, and P_right_ is located about 250 bp from the left end. If these are the terminal promoters, then it is not expected that the regions upstream of P_left_ and P_right_ would be transcribed. Surprisingly, we observed expression from this region. A contig of 1169 bp was identified spanning the cos site, starting at 12 bp on the left arm, crossing over the cos site, and ending at nt position 47843, which is 1283 bp from the left end. There is a discrepancy between the observed size of 1169 (NC1 segments 1, 2, and 3) and the size obtained from the map, which is 1295 bp (1283 bp + 12 bp). This discrepancy is due to a region of about 100 bp that we found to be deleted in the D29 strain we used compared to the wild-type strain. We also observed a transcribed region, NC5, originating from the left end. How these transcripts originate is an intriguing question. Interestingly, a part of the NC segment 2 region is highly expressed and corresponds to the non-coding RNA region of L5 that was deleted in the Δ2 mutant reported earlier (15). The expression of regions upstream of P_left_ and P_righ_t remains a mystery. Still, one definitive point emerges: the left and right arms combine following infection, forming a closed circle or concatamers, and the junction region is also transcribed —an observation not made in any of the earlier studies.

We were curious to see how both the phage and host transcriptomes change as the growth temperature of D29ts12 is increased from 37 to 42 °C. We found that left-arm gene expression increased manifold at the non-permissive temperature of 42°C, whereas right-arm gene expression was completely shut down. The altered pattern persisted regardless of time (30 or 120 minutes), although the magnitude of change was significantly greater at 120 minutes than at 30 minutes. The results indicate that the hypothetical temperature-sensitive factor in D29ts12 controls expression from the two arms in opposing ways. Whereas it appears to influence the right-arm genes positively, it has the opposite effect on the left-arm genes. Thus, there must be a mechanism that balances the expression from the two arms. Abnormally high expression from the left arm may cause early lysis of the host cells, and therefore, expression from this arm must be prevented.

We then compared the effect of phage growth on host gene expression. We argued that at 37°C, where phage growth is permissible, host gene expression is likely to be affected, as seen with lytic phages such as T4 (26). On the other hand, since at 42°C, the phage ceases to grow, the alteration in host gene expression would be minimised. By comparing the transcriptomes, we identified several genes that were differentially expressed. The proteins encoded by these genes were then subjected to KEGG analysis. The analysis revealed an intriguing insight into the nature of the proteins encoded by the differentially regulated genes. One of the conclusions derived was that the most affected group of proteins was ABC transporters. We could group these transporters based on their proposed functions.

The first group of ABC transporters is thought to be involved in Fe transport. The genes are MSMEG_0012, MSMEG_0015, MSMEG_0018 and MSMEG_0020. Iron is an important component of mycobacterial physiology. Many components of mycobacteria, such as cytochromes, catalase, peroxidase, and iron-sulfur clusters, contain iron. TB patients tend to be anaemic. However, this anaemic condition appears to be beneficial as in such situations the viability of Mtb is severely compromised (27). In mycobacteria, iron uptake occurs via two siderophores: mycobactin and exochelin (28, 29). The mycobactins (MBs) are derivatives of the phenyloxazolidine ring, whereas the exochelins are peptidic siderophores whose iron-chelating ability is associated with ornithine-derived hydroxymates. MBs and exochelins are complex molecules with high affinity for iron and are integral to the bacterial strategy for acquiring iron from the environment. Mycobacteria may use either MBs, exocins, or both for iron uptake. Thus, Mtb uses only MBs as siderophores*, M. vaccae* uses only exochelins, whereas *M. smegmatis* uses both MB and exochelin for iron acquisition. Exochelin biosynthesis involves four genes: fxuA, fxuB, fxuC, and *fxbA.* Some of these genes are homologs of *E. coli* genes involved in the production of siderophores known as enterobactins (29). The functions of the genes present in the iron acquisition cluster are as follows: MSMEG_0011, Iron utilisation protein; MSMEG_0012, Ferric exochelin uptake (FxuC); MSMEG_0013, Ferric exochelin uptake (FxuA); MSMEG_0014, Ferric exochelin uptake (FxuB); MSMEG_0015, Ferric exochelin biosynthesis (FxbA); MSMEG_0016, MbtH protein–related protein; MSMEG_0017, Exit protein; MSMEG_0018, Exit protein; MSMEG_0019, Peptide synthetase homolog; MSMEG_0020, FxuD protein. All these genes are clustered and are likely regulated by IdeR, the regulator of genes involved in iron uptake (30). In our transcriptomic studies, we detected transcripts corresponding to FxuC, FxbA, FxuD, and another transporter. Our results show that in all these cases, significant downregulation was observed. FxbA encodes a putative formyltransferase, an essential enzyme in the exochelin synthesis pathway. Therefore, a reduction in FxbA levels should impair exochelin production under iron-limited conditions (29).

Three other gene sets were differentially regulated: one related to the sugar transport system was upregulated, and another group, comprising a pair of genes encoding the phosphate transfer system, was downregulated. Another gene that was downregulated was MSMEG_6880, which encodes a putative hydrophobic amino acid transporter. MSMEG_6880 belongs to a cluster comprising MSMEG_6879, MSMEG_6878, and MSMEG_6877, all of which are linked to neutral amino acid transport. This entire cluster appears to be dedicated to transporting hydrophobic amino acids, which include branched-chain amino acids. Branched-chain amino acids play an essential role in starvation signalling in Gram-positive bacteria; therefore, the decay of transcripts encoding the transporters responsible for their uptake is likely to have a significant impact on the viability of mycobacteria (31, 32). It is interesting to note that genes encoding enzymes required for branched-chain amino acid synthesis are also downregulated.

There is an apparent discrepancy between the RNA-seq and RT-PCR results. The differential expression pattern obtained from direct D29 expression appears to be the opposite of that observed in RNA-seq experiments. Thus, for example, at the non-permissive temperature, expression of several genes decreased, indicating that, in the wild-type infection, these genes are expected to increase; however, in the RT-PCR experiment, we observed a decrease. We can explain this by noting that, unlike initially thought, the temperature shift did not result in a complete shutdown of gene expression. The left arm was not only transcriptionally active, but increasingly so. Therefore, there is always the possibility that the residual transcriptional activity from the mutant phage genome after thermal inactivation is sufficient to alter host gene expression in the same way as the wild type does.

To unravel the genetic mechanism underlying phage-induced mycobacterial PCD, we previously investigated the role of one of its key regulators, the Toxin-Antitoxin (TA) modules. Msm has three TA systems: VapBC, MazEF, and phd/doc(19). We compared the RNA levels of the toxins and antitoxin genes in Msm uninfected and infected cells. We found that the VapB RNA content was reduced relative to VapC. In other words, there was a relative increase in the level of VapC (8).

In a previous study, VapC and VapBC were overexpressed in a VapBC-minus Msm mutant, and the expression levels of individual genes under VapC relative to VapBC were measured. It may be mentioned that whereas VapC is a toxin, the VapBC complex is not. The results of such an experiment indicated that a set of 42 genes was downregulated in VapC-expressing cells compared with VapBC (19). We found that following phage infection, all but six genes were significantly down-regulated. This result supports our contention that phage infection may be increasing VapC activity. Of the six genes that were not down-regulated, three were substantially upregulated. These were MSMEG_3264, 3265 and 3267. Of these three genes, the first, MSMEG_3264, has been predicted to be a transcriptional regulator. MSMEG_3265 is a possible sorbitol dehydrogenase, and 3267 is an erythritol transporter. These three genes may be involved in the transport of sugar alcohols. Why these three genes behaved differently is an intriguing issue. It may be that the expression of these genes is necessary for phage infection, and that an unknown mechanism therefore protects their transcripts.

We verified most of the NGS results from the experiment using D29-infected cells by performing RT-PCR on a subset of genes. Specifically, we focused on a set of genes encoding enzymes involved in glycerol metabolism. The results indicate that the expression of all these genes was downregulated. All these genes belong to the group of ‘42’ that were found to be downregulated following overexpression of VapC as a reporter in the previous study.

In addition to glycerol metabolism genes, we also targeted two marker genes, MSMEG_1712 and MSMEG_4643. In the paper on the effect of VapC expression on carbon metabolism (19), MSMEG_1712 was reported to be one of the genes highly downregulated in VapC-expressing cells compared with VapBC-expressing cells. In contrast, another gene, MSMEG_4643, was moderately upregulated. A significant downregulation of MSMEG_1712 and a moderate increase in the expression of MSMEG_4643 appear to be the signature of VapC’s activity. We argued that if phage D29 were activating VapC, then the level of MSMEG_1712 transcripts should decrease, and that of MSMEG_4643 must either increase or at least remain the same. The results were consistent with such a hypothesis. All these observations indicate that phage D29 may be activating VapC.

In this study, we assessed for the first time the expression patterns of phage D29 genes and host genes. The information presented will help further our understanding of how D29 inactivates its host following infection.

## Acknowledgments

We thank Dr R. McNerney for the D29 phage. R.S. acknowledge the University Grants Commission of India (UGC India), Govt. of India, for their financial aid. SKDG acknowledges receiving support from CSIR in the form of an Emeritus fellowship (Sanction no. 21/(1134)/22/EMR II) dated 19.05.2022. SKDG acknowledges the assistance of Ms Byapti Ghosh and Dr Zhumur Ghosh, of the Bioinformatics division of Bose Institute, for interpreting and analysing the NGS data.

## Notes

### Competing Interest Statement

The authors have declared no competing interest.

## References

1. McNerney R. TB: the return of the phage. A review of fifty years of mycobacteriophage research. Int J Tuberc Lung Dis. 1999;3(3):179–84.

2. Pope WH, Hatfull GF. Adding pieces to the puzzle: New insights into bacteriophage diversity from integrated research-education programs. Bacteriophage. 2015;5(4):e1084073.

3. Guerrero-Bustamante CA, Dedrick RM, Garlena RA, Russell DA, Hatfull GF. Toward a Phage Cocktail for Tuberculosis: Susceptibility and Tuberculocidal Action of Mycobacteriophages against Diverse Mycobacterium tuberculosis Strains. mBio. 2021;12(3).

4. Lopatina A, Tal N, Sorek R. Abortive Infection: Bacterial Suicide as an Antiviral Immune Strategy. Annu Rev Virol. 2020;7(1):371–84.

5. Jenson JM, Chen ZJ. cGAS goes viral: A conserved immune defense system from bacteria to humans. Mol Cell. 2024;84(1):120–30.

6. Samaddar S, Grewal RK, Sinha S, Ghosh S, Roy S, Das Gupta SK. Dynamics of Mycobacteriophage-Mycobacterial Host Interaction: Evidence for Secondary Mechanisms for Host Lethality. Appl Environ Microbiol. 2016;82(1):124–33.

7. Allocati N, Masulli M, Di Ilio C, De Laurenzi V. Die for the community: an overview of programmed cell death in bacteria. Cell Death Dis. 2015;6(1):e1609.

8. Calcuttawala F, Shaw R, Sarbajna A, Dutta M, Sinha S, S KDG. Apoptosis like symptoms associated with abortive infection of Mycobacterium smegmatis by mycobacteriophage D29. PLoS One. 2022;17(5):e0259480.

9. Tokunaga T, Sellers M. Infection of Mycobacterium Smegmatis with D29 Phage DNA. J Exp Med. 1964;119:139–49.

10. Grange JM. The genetics of mycobacteria and mycobacteriophages - a review. Tubercle. 1975;56(3):227–38.

11. Ford ME, Sarkis GJ, Belanger AE, Hendrix RW, Hatfull GF. Genome structure of mycobacteriophage D29: implications for phage evolution. J Mol Biol. 1998;279(1):143–64.

12. Bhattacharya B, Giri N, Mitra M, Gupta SK. Cloning, characterization and expression analysis of nucleotide metabolism-related genes of mycobacteriophage L5. FEMS Microbiol Lett. 2008;280(1):64–72.

13. Bavda VR, Yadav A, Jain V. Decoding the molecular properties of mycobacteriophage D29 Holin provides insights into Holin engineering. J Virol. 2021.

14. Oyaski M, Hatfull GF. The cohesive ends of mycobacteriophage L5 DNA. Nucleic Acids Res. 1992;20(12):3251.

15. Dedrick RM, Mavrich TN, Ng WL, Hatfull GF. Expression and evolutionary patterns of mycobacteriophage D29 and its temperate close relatives. BMC Microbiol. 2017;17(1):225.

16. Barman A, Shaw R, Bhawsinghka N, Das Gupta SK. A CRISPRi-based investigation reveals that multiple promoter elements drive gene expression from the genome of mycobacteriophage D29. Microbiology (Reading). 2022;168(11).

17. Bhawsinghka N, Dutta A, Mukhopadhyay J, Gupta SKD. A transcriptomic analysis of the mycobacteriophage D29 genome reveals the presence of novel stoperator-associated promoters in its right arm. Microbiology. 2018;164(9):1168–79.

18. Ghosh S, Shaw R, Sarkar A, Gupta SKD. Evidence of positive regulation of mycobacteriophage D29 early gene expression obtained from an investigation using a temperature-sensitive mutant of the phage. FEMS Microbiol Lett. 2020;367(21).

19. McKenzie JL, Robson J, Berney M, Smith TC, Ruthe A, Gardner PP, et al. A VapBC toxin-antitoxin module is a posttranscriptional regulator of metabolic flux in mycobacteria. J Bacteriol. 2012;194(9):2189–204.

20. Bhawsinghka N, Dutta A, Mukhopadhyay J, Das Gupta SK. A transcriptomic analysis of the mycobacteriophage D29 genome reveals the presence of novel stoperator-associated promoters in its right arm. Microbiology. 2018;164(9):1168–79.

21. Langmead B, Salzberg SL. Fast gapped-read alignment with Bowtie 2. Nat Methods. 2012;9(4):357–9.

22. Prjibelski A, Antipov D, Meleshko D, Lapidus A, Korobeynikov A. Using SPAdes De Novo Assembler. Curr Protoc Bioinformatics. 2020;70(1):e102.

23. Li H, Handsaker B, Wysoker A, Fennell T, Ruan J, Homer N, et al. The Sequence Alignment/Map format and SAMtools. Bioinformatics. 2009;25(16):2078–9.

24. Calcuttawala F, Shaw R, Sarbajna A, Dutta M, Sinha S, K. Das Gupta S. Apoptosis like symptoms associated with abortive infection of Mycobacterium smegmatis by mycobacteriophage D29. Plos one. 2022;17(5):e0259480.

25. Bhawsinghka N, Dutta A, Mukhopadhyay J, Das Gupta SK. A transcriptomic analysis of the mycobacteriophage D29 genome reveals the presence of novel stoperator-associated promoters in its right arm. Microbiology (Reading). 2018;164(9):1168–79.

26. Hinton DM. Transcription from a bacteriophage T4 middle promoter using T4 motA protein and phage-modified RNA polymerase. J Biol Chem. 1991;266(27):18034–44.

27. Ratledge C. Iron, mycobacteria and tuberculosis. Tuberculosis (Edinb). 2004;84(1-2):110–30.

28. De Voss JJ, Rutter K, Schroeder BG, Barry CE, 3rd. Iron acquisition and metabolism by mycobacteria. J Bacteriol. 1999;181(15):4443–51.

29. Fiss EH, Yu S, Jacobs WR, Jr. Identification of genes involved in the sequestration of iron in mycobacteria: the ferric exochelin biosynthetic and uptake pathways. Mol Microbiol. 1994;14(3):557–69.

30. Yellaboina S, Ranjan S, Vindal V, Ranjan A. Comparative analysis of iron-regulated genes in mycobacteria. FEBS Lett. 2006;580(11):2567–76.

31. Kaiser JC, Heinrichs DE. Branching Out: Alterations in Bacterial Physiology and Virulence Due to Branched-Chain Amino Acid Deprivation. mBio. 2018;9(5).

32. Guedon E, Serror P, Ehrlich SD, Renault P, Delorme C. Pleiotropic transcriptional repressor CodY senses the intracellular pool of branched-chain amino acids in Lactococcus lactis. Mol Microbiol. 2001;40(5):1227–39.

