## Supplemental figures and tables for "Integrated host–phage RNA seq analysis of Mycobacteriophage D29 infection reveals dual-arm balanced phage genome expression and inhibited expression of host transporter and VapC target genes"

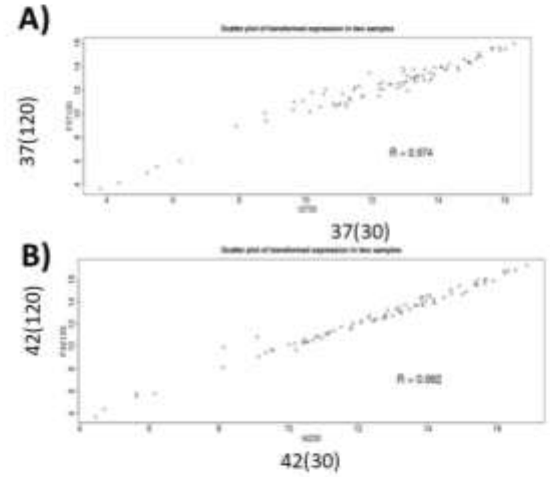

**Fig. S1**. Correlation between the RNA seq results obtained using the samples 37◦C(30 min) and 37◦C(120), (A) and 42◦C(30) and 42◦C(120) (B). In both cases, the R values are close to 1, indicating significant correlation.

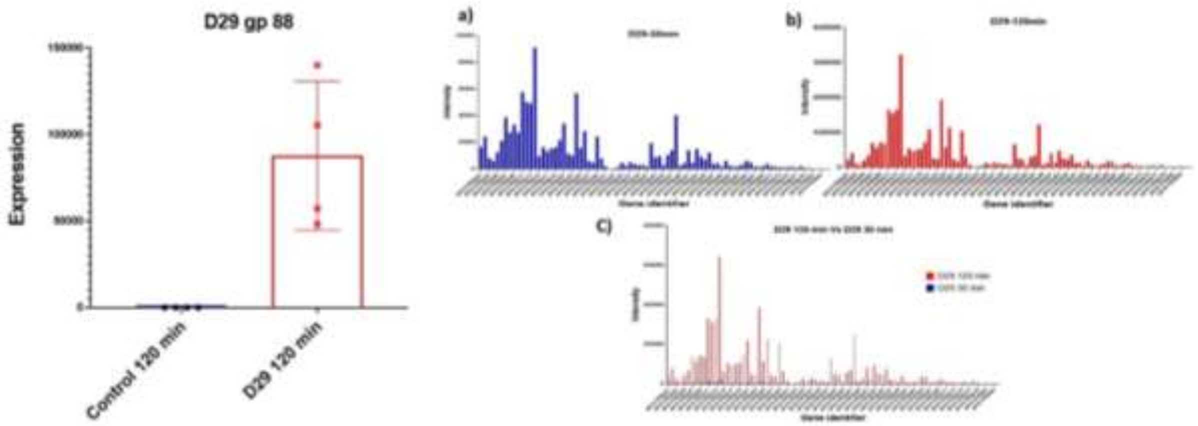

**Fig. S2.** Evidence that Msm cells received phage D29 infection and that the genome was expressed. Left panel, RT-PCR assay for D29 gp88 gene expression a marker gene used in our lab to monitor phage infection and expression. Right panel, Normalised (TPN) gene expression values at the early stage of infection (30 mins.) and late (120 mins) (a and b). In (c) the results of a and b are combined. The red bars represent 120 mins and the relatively small bars (blue) 30 mins. Post-infection.

**Table S1**. Gene ontology enrichment using KEGG pathways for the differentially regulated genes identified through NGS analysis.

*As separate Excel file.*

**Table S2**. Comparative analysis of genes that are differentially expressed in VapC overexpressing cells and phage infected.

| Predicted carbon substrate (s) | Gene ID or locus tag | Description | Control 120min | D29_120min | Fold change 120 min (control 120min / d29 120 min) | Expression Upon D29 infection (Down/Up regulated) | Fold change in response to VapC * | Expression after VapC Over expression (Down/Up) * |
|---|---|---|---|---|---|---|---|---|
| Fructose | MSMEG_0084 | Phosphocarrier protein hpr | 370.3573897 | 139.0368514 | 2.663735448 | Down | 1.28 | Down |
| Glycerol | MSMEG_1542 | Transcriptional regulator | 956.8132197 | 590.1341916 | 1.621348556 | Down | 1.23 | Down |
| | MSMEG_1543 | EPTC-inducible aldehyde dehydrogenase | 39535.14169 | 5261.772399 | 7.513654846 | Down | **2.18** | Down |
| | MSMEG_1546 | Coenzyme B12dependent glycerol dehydrogenase small subunit | 3146.338925 | 701.3636727 | 4.48603064 | Down | **1.98** | Down |
| | MSMEG_1547 | Glycerol dehydratase large subunit | 16567.54708 | 3661.303754 | 4.525040313 | Down | **2.02** | Down |
| Xylose | MSMEG_1704 | ABC transporter | 3793.275136 | 698.2739649 | 5.432359399 | Down | **1.63** | Down |
| Arabinose | MSMEG_1707 | Phosphatase YfbT | 820.2226961 | 278.0737028 | 2.949659345 | Down | 1.23 | Down |
| | MSMEG_1708 | Ribose operon repressor, putative | 580.3398364 | 163.7545139 | 3.543962377 | Down | 1 | Down |

| | | | | | | | | |
|---|---|---|---|---|---|---|---|---|
| | MSMEG_1709 | Inner membrane ABC transporter permease protein YjfF | 891.5759547 | 219.3692545 | 4.064270341 | Down | 1.07 | Down |
| | MSMEG_1714 | l-Ribulose-5-phosphate 4epimerase UlaF | 2044.780524 | 460.3664636 | 4.441636578 | Down | 1.3 | Down |
| | MSMEG_1715 | l-Ribulose-5-phosphate 4epimerase UlaF | 3921.031447 | 1059.769779 | 3.69988985 | Down | 1.48 | Down |
| Glucose, trehalose, *N* acetylglucosamine | MSMEG_2114 | Glucose-6phosphate isomerase, putative | 1741.699064 | 1340.933189 | 1.298870874 | Down | 1.25 | Down |
| | MSMEG_2115 | Conserved hypothetical protein | 14439.86087 | 11660.55727 | 1.238350838 | Down | 1.11 | Down |
| | MSMEG_2117 | b-Glucosidespecific EII permease | 297.6450215 | 108.1397733 | 2.752410258 | Down | 1.32 | Down |
| | MSMEG_2118 | Glucosamine-6phosphate isomerase | 625.8700109 | 203.9207154 | 3.069183087 | Down | 1.37 | Down |
| | MSMEG_2119 | *N*-Acetylglucosamine-6-phosphate deacetylase | 971.0838714 | 423.2899699 | 2.294133905 | Down | 1.38 | Down |
| Dihydroxyacetone | MSMEG_2122 | Dihydroxyacetone kinase, L subunit | 4467.393541 | 812.5931539 | 5.497700196 | Down | **1.96** | Down |
| | MSMEG_2124 | Glycerol uptake facilitator, MIP channel | 1660.832038 | 494.3532495 | 3.359605786 | Down | **2.73** | Down |
| Ribose, ribonucleosides | MSMEG_3092 | Transcriptional regulator, sugar-binding family protein | 184.1593626 | 163.7545139 | 1.124606328 | Down | 2.08 | Down |
| Unknown | MSMEG_3264 | Transcriptional regulator | 120.9607621 | 599.403315 | 0.201801957 | Up | 1.07 | Down |

| | | | | | | | | |
|---|---|---|---|---|---|---|---|---|
| | MSMEG_3265 | Arabitol-phosphate dehydrogenase | 484.5226034 | 2335.819104 | 0.207431561 | Up | 1.38 | Down |
| | MSMEG_3267 | Transporter | 159.6953882 | 1170.99926 | 0.136375311 | Up | **1.69** | Down |
| Ribose | MSMEG_3598 | Periplasmic sugarbinding proteins | 1353.673248 | 1059.769779 | 1.277327657 | Down | 1.28 | Down |
| | MSMEG_3599 | Sugar-binding transcriptional regulator, LacI family protein | 1319.695506 | 892.9255569 | 1.477945721 | Down | 1.42 | Down |
| | MSMEG_3601 | Ribose/xylose/arabinose/galactoside ABC-type transport systems, permease components | 347.9320799 | 441.8282167 | 0.787482706 | Up | **1.53** | Down |
| | MSMEG_3602 | Ribose transport ATP-binding protein RbsA | 455.9813 | 537.6091588 | 0.848165052 | Up | 1.22 | Down |
| Unknown sugar | MSMEG_4657 | ABC transporter membrane protein | 771.2947474 | 247.1766247 | 3.12041945 | Down | 1.42 | Down |
| | MSMEG_4658 | Sugar ABC transporter substrate-binding protein | 997.5865103 | 466.5458792 | 2.13823882 | Down | 1.49 | Down |
| | MSMEG_4659 | GntR family protein transcriptional regulator | 254.1535115 | 262.6251638 | 0.96774242 | Up | 1.02 | Down |
| | MSMEG_4662 | Deoxyribosephosphate aldolase superfamily protein | 170.5682657 | 160.6648061 | 1.061640504 | Down | 1.27 | Down |
| | MSMEG_5057 | Conserved hypothetical protein | 1202.812073 | 577.7753604 | 2.081798837 | Down | 1.36 | Down |
| | MSMEG_5060 | ABC transporter, permease protein SugA | 1386.291881 | 713.722504 | 1.942340157 | Down | 1.37 | Down |
| | MSMEG_5062 | Conserved hypothetical protein | 327.5454346 | 176.1133451 | 1.859855846 | Down | 1.11 | Down |

| | | | | | | | | |
|---|---|---|---|---|---|---|---|---|
| Succinate | MSMEG _5302 | Aerobic $C_4$dicarboxylate transport protein | 2385.917056 | 1489.239164 | 1.602104694 | Down | **1.73** | Down |
| | MSMEG _5572 | Sugar ABC transporter permease protein | 603.444701 | 160.6648061 | 3.755923377 | Down | **1.76** | Down |
| | MSMEG _5573 | Sugar ABC transporter permease protein | 748.8694375 | 176.1133451 | 4.252201541 | Down | **1.99** | Down |
| Sugar alcohol | MSMEG _5574 | Substrate-binding protein | 1030.884698 | 407.8414308 | 2.527660555 | Down | **1.92** | Down |
| | MSMEG _5575 | Repressor | 1334.645713 | 333.6884434 | 3.999676162 | Down | **1.68** | Down |
| | MSMEG _6019 | ABC-type sugar transport system ATPase component | 1189.220976 | 188.4721764 | 6.309795955 | Down | 1.2 | Down |
| Xylose | MSMEG _6020 | d-Xylose-binding periplasmic protein | 1680.539128 | 240.9972091 | 6.973272156 | Down | 1.11 | Down |
| | MSMEG _6021 | Xylose isomerase | 732.5601213 | 132.8574358 | 5.513881228 | Down | **1.65** | Down |
| Glycerol | MSMEG _6239 | 1,3-Propanediol dehydrogenase | 2971.013776 | 528.3400354 | 5.623298589 | Down | **1.77** | Down |

* The data presented in this column has been taken from Ref.  54. with permission.
